# Interplay between disordered regions in RNAs and proteins modulates interactions within stress granules and processing bodies

**DOI:** 10.1101/2021.05.05.442738

**Authors:** Andrea Vandelli, Fernando Cid Samper, Marc Torrent Burgas, Natalia Sanchez de Groot, Gian Gaetano Tartaglia

**Affiliations:** Department of Biochemistry and Molecular Biology, Universitat Autònoma de Barcelona, Bellaterra, 08193, Barcelona, Spain; Universitat Pompeu Fabra (UPF), 08003 Barcelona, Spain; Centre for Genomic Regulation (CRG), The Barcelona Institute for Science and Technology, 08003, Barcelona, Spain; Center for Human Technologies, Istituto Italiano di Tecnologia,16152, Genova, Italy; Department of Biology ‘Charles Darwin’, Sapienza University of Rome, 00185, Rome, Italy; Institucio Catalana de Recerca i Estudis Avançats (ICREA), 08010, Barcelona, Spain

**Keywords:** Structural disorder, interaction networks, RNA-binding protein, stress granule, processing bodies, RNA structure, biological condensates, liquid-liquid phase separation, fuzzy interactions

## Abstract

Condensation, or liquid-like phase separation, is a phenomenon indispensable for the spatiotemporal regulation of molecules within the cell. Recent studies indicate that the composition and molecular organization of phase-separated organelles such as Stress Granules (SGs) and Processing Bodies (PBs) are highly variable and dynamic. A dense contact network involving both RNAs and proteins controls the formation of SGs and PBs and an intricate molecular architecture, at present poorly understood, guarantees that these assemblies sense and adapt to different stresses and environmental changes. Here, we investigated the physico-chemical properties of SGs and PBs components and studied the architecture of their interaction networks. We found that proteins and RNAs establishing the largest amount of contacts in SGs and PBs have distinct structural properties and intrinsic disorder is enriched in all protein-RNA, protein-protein and RNA-RNA interaction networks. The increase of disorder in proteins is accompanied by an enrichment in single-stranded regions of RNA binding partners. Our results suggest that SGs and PBs quickly assemble and disassemble through fuzzy-like dynamic contacts modulated by unfolded domains of their components.

**Research Highlights:**

- We systematically studied RNA-RNA, protein-protein and RNA-protein interaction networks in stress granules and processing bodies;
- RNAs enriched in stress granules and processing bodies are more single-stranded and form a large number of contacts with both proteins and RNAs;
- Proteins in stress granules and processing bodies are less structured and contact larger amounts of single-stranded RNAs.

## INTRODUCTION

Cells exploit the spatiotemporal confinement for efficient organization of biochemical reactions [1]. In the complex and crowded intracellular milieu [2], condensation in membrane-bound or membrane-less organelles allows to control concentration and interactions of the reactants [3]. These assemblies, located in both cytoplasm and nucleus, participate in multiple cellular functions [4] including stress response, transport channels in the nuclear pore complex and chromatin reorganization [5].

Molecular condensation is currently the subject of intense investigation and recent advances started to reveal their composition and inner architecture [3,6,7]. Molecular interactions within molecular condensates are not yet understood, but involve proteins and RNAs [8,9]. These assemblies have liquid-like properties and are commonly formed through a process that requires phase separation [10,11]. Valency or number of interaction sites dictates the contact density of the molecular network regulating the stability, organization and composition of the condensates [10,12].

Through helical interactions, canonical and non-canonical Watson-Crick base-pairing, RNAs interact with other RNAs and promote phase separation [13]. Yet, technical difficulties in the study of RNA-RNA contacts currently impede our complete understanding of this phenomenon [14]. Protein-protein interactions, and especially prion-like elements, contribute to condensation by promoting protein associations [10,15]. More specifically, perturbation of the native state [16] accompanied by an increase in structural disorder [17] and hydrophobicity [18] enhance the propensity of proteins to aggregate [15].

Depending on their binding preferences, RNA-binding proteins (RBPs) interact with either single or double-stranded regions of RNAs [19]. Highly structured RNAs attract large amounts of proteins thanks to their intrinsic ability to establish stable interactions [12,20]. RNAs are scaffolding elements: whereas a polypeptide of 100 amino acids can interact with one or two proteins, a chain of 100 nucleotides is able to bind to 5-20 proteins [21]. Not only RNA attracts proteins, but also proteins can in turn contribute to change RNA properties: chemical modifications such as N^1^-methyladenosine (m1A) and N^6^-methyladenosine (m6A) can modify RNA structure [22,23] and influence the formation of ribonucleoprotein condensates [24,25]. Helicases such as the Eukaryotic initiation factor 4A-I can also alter RNA structure by opening up double-stranded regions and altering cellular interactions [26].

Here, we used a computational approach to investigate the interactions and properties of RNA and protein in the two of the best-known biological condensates: stress granules (SGs) [27] and processing-bodies (PBs) [28]. These large assemblies arise upon viral infection or when chemical and physical insults occur to cells. They are thought to form to protect transcripts that would otherwise be aberrantly processed. More specifically, SGs store non-translating mRNAs as indicated by translation initiation factors enriched in the pool of proteins that compose them, whereas PBs facilitate RNA decay because of the abundance in RNA decapping and deadenylation enzymes [29].

Proteins [7,9] and RNAs [27,30] contained in SGs and PBs are only now starting to be unveiled and their interaction networks are largely unknown. With the present systematic analysis, we aim to characterize how structure influences the interactions sustaining these biological condensates, including both proteins and RNAs and all its possible combinations (RNA-RNA, protein-protein and RNA-protein). Our results show similarities and interconnections between the most contacted players from both molecular types. We report the intriguing result that RNAs enriched in SGs and PBs are more disordered and form a larger number of contacts with RNAs and proteins. At the same time, proteins enriched in SGs and PBs are more disordered and form a larger number of contacts with proteins and RNAs. Taken together, our data suggest that structural disorder is a property that distinguishes dynamic fuzzy-like assemblies such as PBs and SGs from other solid-like aggregates [31,32].

## RESULTS

### RNA structure drives interaction with proteins in SGs and PBs

SGs and PBs are two of the best-known biological condensates. They contain multiple proteins whose concentration changes with stress, cell state and environmental conditions [9,29]. Between them it has been found a small specific set of proteins essential for their formation involved in the recruitment of the other components and in sustaining the condensate. Despite this, there are still uncertainties regarding how the cell regulates their content and assembly.

We recently reported that protein-RNA interactions build up the scaffold of phase-separating organelles [10,33] and their selective recruitment is dictated by RNA physicochemical properties [12,20]. Specifically, we have shown that RNAs engaging in interactions with many protein partners are enriched in double-stranded content (**Figure 1A**) [19,20]. The origin of this property, observed with a number of different experimental approaches, is that double-stranded regions reduce the intrinsic flexibility of the polynucleotide chain. Presence of a stable fold favors the formation of stable and well-defined binding sites where the protein can bind. However, our observation does not suggest that protein binding sites and double□stranded regions are the same. If a specific interaction occurs in a small loop at the end of a stem, the overall region is enriched in double□stranded nucleotides, although the exact binding could be in a single□stranded region.

**Figure 1.**
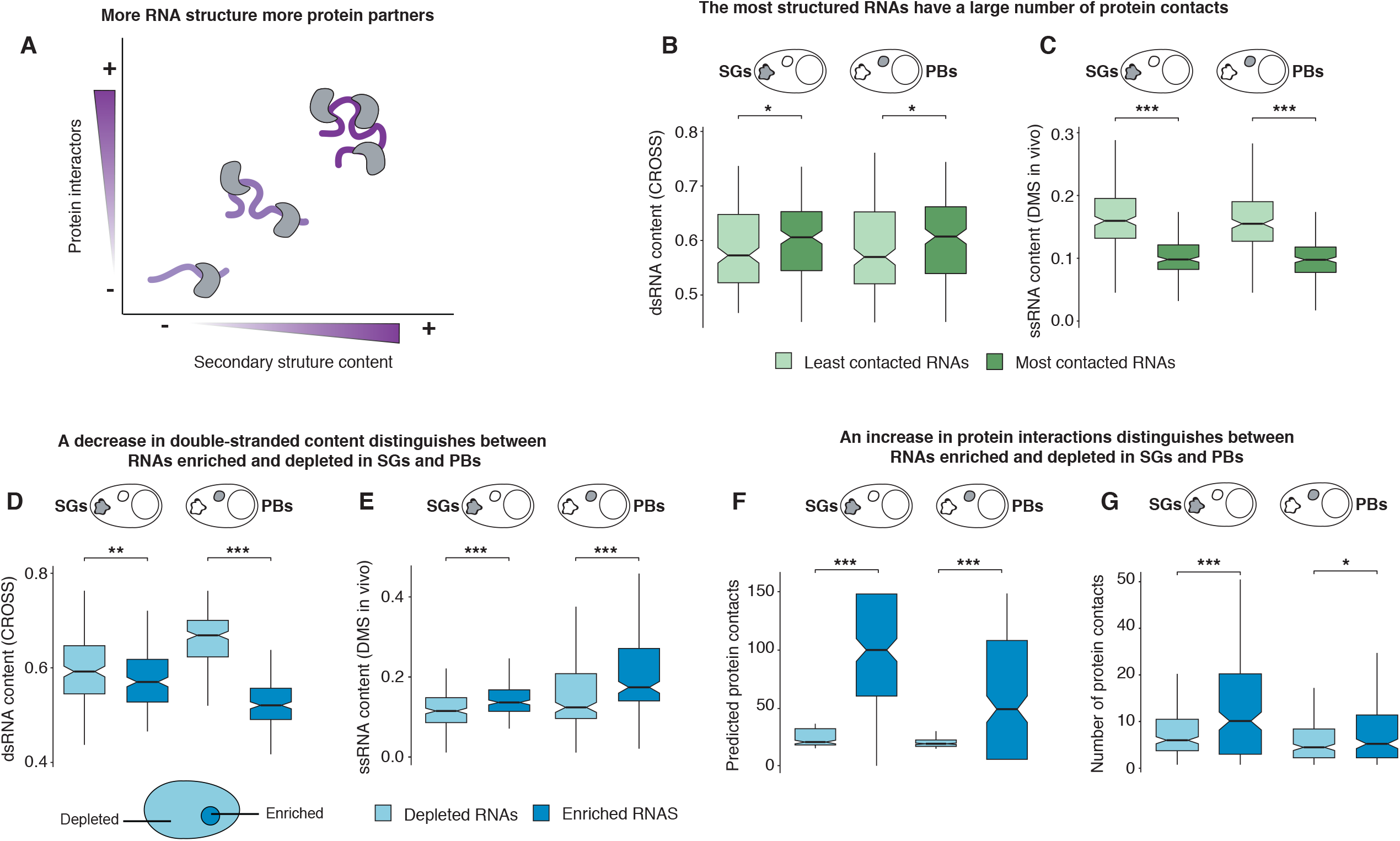
RNAs enriched in SGs and PBs are less structured and contact a larger amount of proteins. **A**. Graphical representation of the relationship between number of protein interactions and double-stranded content of RNAs. The trend was identified by using different computational and experimental techniques; **B**. Double-stranded content dsRNA (*CROSS* predictions) of RNAs present in SGs and PBs. The RNAs are categorized in two classes: least- and most-contacted depending on the amount of protein interactions detected by eCLIP. An equal amount of 200 transcripts is used in each class (SGs and PBs, least and most contacted RNAs). Significant differentiation is found (SG p-value < 0.013 and PB p-value < 0.029, Wilcoxon test); **C**. Single stranded content ssRNA (dimethyl sulfate modification, DMS, measured *in vivo*) for RNAs most and least contacted by proteins in SGs and PBs. RNA classes follow the definition given in panel **B**. Significant differentiation is found (SG p-value <1.15e-34, PB p-value < 3.19e-35, Wilcoxon test); **D**. Double stranded content dsRNA (CROSS predictions) for RNAs enriched or depleted in SGs and PBs. An equal amount of 200 transcripts is used for each category (SGs and PBs, depleted and enriched RNAs). Significant differentiation is found (SG p-value < 0.006, PB: p-value < 1.88e-54, Wilcoxon test); **E**. Single stranded content ssRNA (DMS, measured *in vivo*) for RNAs enriched or depleted in SGs and PBs. RNA classes follow the definition given in panel **D**. Significant differentiation is found (SG p-value < 4.51e-67, PB p-value < 2.43e-67, Wilcoxon test); **F**. *cat*RAPID predictions of protein interactions with RNAs enriched or depleted in SGs and PBs. RNA classes follow the definition given in panel **D**. Significant differentiation is found (SG p-value < 3.69e-33, PB p-value < 4.62e-21, Wilcoxon test); **G**. eCLIP detection of protein interactions with RNAs enriched or depleted in SGs and PBs. RNA classes follow the definition of panel **D**. Significant differentiation is found (SG p-value < 1.44e-06, PB p-value < 0.075, Wilcoxon test). Significance indicated in the plots: ^*^ p-value < 0.1, ^**^ p-value < 0.01 and ^***^ p-value < 0.001.

We wondered whether RNA structure drives the interaction with proteins present in SGs and PBs as detected in the whole transcriptome analysis [19,20]. Following up on our previous computational analysis [19,20], we used protein-RNA interactions available from enhanced CLIP (eCLIP) experiments [34] to rank protein associations with RNAs present in SGs and PBs (**Materials and Methods**). We first selected the transcripts with the largest and lowest amount of protein contacts from the list of RNA reported in SGs [27] and PBs [30] (**Supplementary Table 1**) and then compared their secondary structure content. We used *CROSS* [35] to predict the secondary structure properties of transcripts using the information contained in their sequences and we found that RNAs with more protein contacts in SGs and PBs are significantly more structured (**Figure 1B; Materials and Methods**).

*CROSS* reproduces transcriptomic experiments such as *in vivo* click selective 2-hydroxyl acylation and profiling experiment (icSHAPE) [22] and Parallel Analysis of RNA Structure (PARS) [36] with accuracies higher than 0.80 [37]. To assess whether the calculations are in agreement with experimental data, we used data coming from dimethyl sulfate (DMS) foot-printing experiments carried out *in vitro* and *in vivo* [38] (**Materials and Methods**). DMS modification of the unpaired adenosine and cytidine nucleotides is commonly used for revealing structural properties of RNA molecules [39]. The results are in complete accordance with *CROSS* predictions, with the most contacted RNAs being more structured than the least contacted ones (**Figure 1C** and **Supplementary Figure 1**). Although the conditions in which DMS experiments were performed did not take into account formation of SGs and PBs, our results show that for both SGs and PBs the amount of double-stranded regions is statistically associated with the number of protein contacts SG’s and PB’s RNAs can form (**Figure 1C**).

Our results indicate that RNAs establishing interactions with a large number of proteins [12,40] act as scaffolds for the formation of ribonucleoprotein complexes [33,41], which suggests that specific transcripts could be the ‘hubs’ in the transcriptional and post-transcriptional layers of regulation[19,20]. This observation indicates that RNAs could be regarded as network connectors or ‘kinetic condensers’ sustaining and capturing the different components of the biological condensates. Actually, recent evidence indicates that RNA interactions with other RNAs occur spontaneously, thus an additional level exists in the inner regulation of SGs and PBs architecture [13].

### RNA enriched in SGs and PBs are less structured

Since SGs can contain mRNAs from essentially every expressed gene [27], we decided to study a subset of RNAs that are specifically enriched in SGs or PBs. Indeed, it is possible to distinguish two subsets of transcripts, *enriched* and *depleted*, depending on their abundance in SGs and PBs relative to the rest of the transcriptome **(Materials and Methods** and **Supplementary Table 1)**. We stress that the distribution in these groups is independent of the total transcript abundance or the AU content [42]. We also note that the overlap between SGs and PBs is just 25%, and this percentage varies when comparing different sets. Despite these differences, enriched RNAs share similar properties in both cases: they are composed by transcripts with less translation efficiency and longer sequences [27,28].

Since longer sequences have higher probability to have a larger number of interaction partners [7,9], we expected to find an enrichment of double-stranded regions in PBs and SGs [20]. However, our predictions carried out with *CROSS* indicate that these RNAs contain more single-stranded regions than depleted transcripts (**Figure 1D**).

To assess whether the calculations are in agreement with experimental data, we compared our predictions with DMS experiments [38]. The results are in complete accordance, with enriched RNAs being more unstructured than depleted RNAs (**Figure 1E**). Interestingly, the 5’ UTRs, CDS and 3’ UTRs consistently show a lower amount of structure, which indicates that the trend identified is particularly robust (**Supplementary Figure 2**). We obtained similar results using another experimental approach to reveal RNA secondary structure, PARS (**Supplementary Figure 3**) [36]. PARS is an approach based on deep sequencing fragments of RNAs treated with structure-specific enzymes [43] (**Materials and Methods**). Again, RNAs enriched in SGs and PBs have a significantly increased number of single-stranded regions.

### RNAs enriched in SGs and PBs bind a large amount of proteins

We next investigated protein interactions with RNAs enriched in SGs and PBs. In this context, we previously showed that the interactions between proteins and RNAs could scaffold the formation of phase-separating organelles [12,21,33] and the incorporation of RNAs depends on their physico-chemical properties [19,20]. We used the *cat*RAPID approach to predict RNA interactions with proteins (**Materials and Methods**) [44,45]. *cat*RAPID exploits secondary structure predictions coupled with hydrogen bonding and van der Waals calculations to estimate the binding affinity of protein-RNA pairs with an average accuracy of 0.78 [46,47]. For both SGs and PBs, our predictions indicate that enriched RNAs have a significantly larger number of interactions with proteins than depleted RNAs (**Figure 1F**).

To experimentally validate our predictions, we retrieved protein-RNA interactions available from eCLIP experiments (**Materials and Methods**) [34]. On the same set of proteins investigated with *cat*RAPID, we observed that RNAs enriched in PBs and SGs have an increased number of protein partners (**Figure 1G**).

Although predictions and experiments used in our analysis do not consider the cellular context in which SGs and PBs are formed, our models are based on physico-chemical properties of the molecules involved and they are therefore expected to have general validity [19,20]. Intriguingly, RNAs enriched in SGs and PBs establish a dense network of contacts with proteins despite their increase in single-stranded content. This contradicts the trend previously identified and suggests that these RNAs might have an interaction network that deviate from those characterizing the average transcriptome [19,20].

### Single stranded regions are involved in RNA-RNA interactions

Even though protein interactions correlate with the amount of double stranded regions found in them (**Figure 1B**), it is possible that other RNA properties are involved in different interactions. We hypothesized that RNAs enriched in SGs and PBs may interact among themselves through a mechanism of base-pairing recognition in single-stranded regions [48]. To investigate if an increase in single-stranded regions is a property favouring contacts among RNAs, we compared the structures of RNAs that build a larger number of contacts with RNAs and those more prone to interact with proteins (**Figure 2A**). The analysis of the DMS structure shows that the RNAs interacting with a larger number of RNAs are more single-stranded.

**Figure 2.**
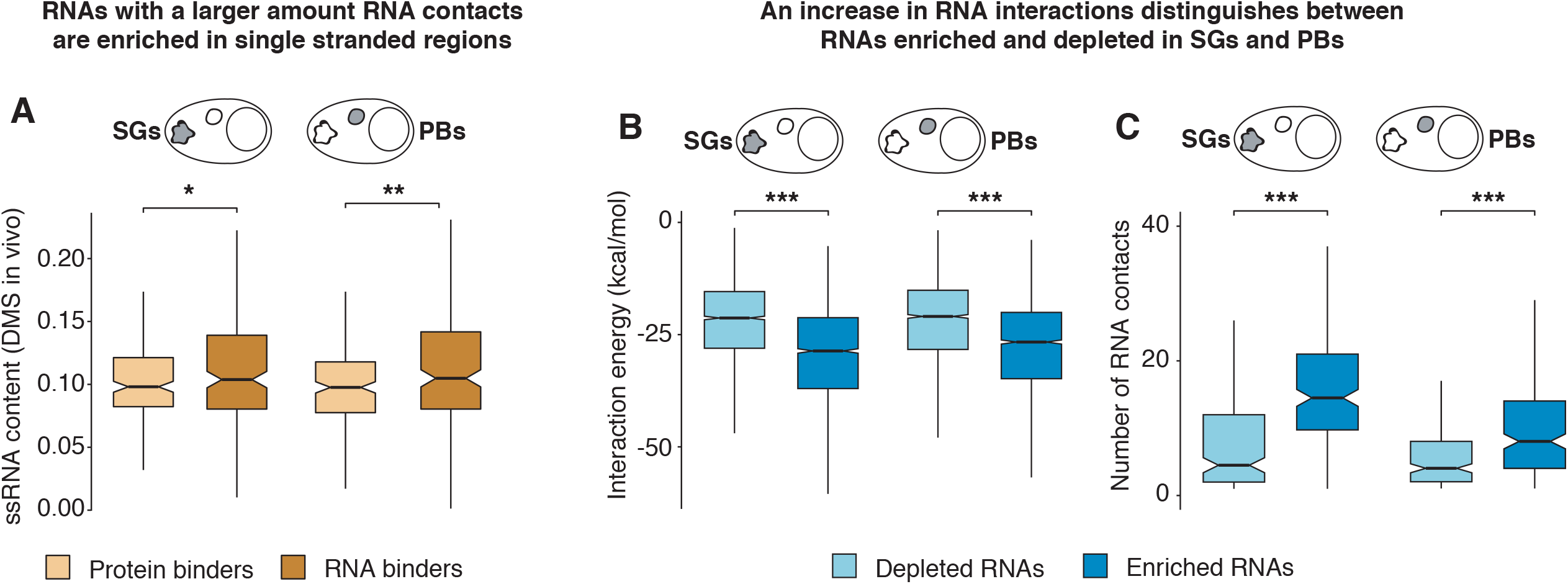
RNAs enriched in SGs and PBs are less structured and contact a larger amount of RNAs. **A**. Single-stranded content ssRNA (DMS measured *in vivo*) of RNAs enriched in protein interactions (eCLIP experiments) and RNAs enriched in RNA interactions (RISE database). An equal amount of 200 transcripts is used in each category (SGs and PBs, protein and RNA binders). Significant differentiation is found (SG p-value < 0.091, PB p-value < 0.007, Wilcoxon test). **B**. IntaRNA predictions of the energies associated with RNA-RNA interactions. An equal amount of 2500 predicted interactions are used in each category (50 transcripts from SGs and PBs, enriched and depleted interacting with 50 random human RNAs). Significant differentiation is found (SG p-value < 2.54e-108, PB p-value < 1.57e-69, Wilcoxon test). For each plot the interaction bonding energy is reported (kcal/mol). **C**. Number of RNA-RNA interactions (RISE database). An equal amount of 200 transcripts is used in each category (SGs and PBs, enriched and depleted). Significant differentiation is found (SG p-value < 1.54e-19, PB p-value < 5.89e-09, Wilcoxon test). Significance indicated in the plots: ^*^ p-value < 0.1, ^**^ p-value < 0.01 and ^***^ p-value < 0.001.

Using IntaRNA to predict RNA-RNA interactions (**Materials and Methods**) [48], we then compared the binding ability of the most enriched and depleted RNAs in SGs and PBs. Our results clearly show that enriched RNAs are more prone to interact with RNAs (**Figure 2B**).

We then searched available experimental data to validate our predictions. To this aim, we used the RISE database containing RNA-RNA interactions assessed through high-throughput approaches [49]. By counting the number of binding partners that each transcript has with other transcripts (**Materials and Methods**), we found that the enriched RNAs are associated with a large number of binding partners (**Figure 2C**). Altogether, our results indicate that enriched RNAs are more single-stranded and base-pair with multiple RNAs to establish a larger number of contacts.

Thus, RNAs enriched in SGs and PBs are able to establish a dense network of contacts not just with proteins but also with RNAs. This result suggests that RNAs in SGs and PBs could act as central players sustaining their inner architecture.

### Enriched RNAs are populated by master regulators of protein- and RNA-binding

To understand how enriched RNAs are able to create a dense network of contacts, we studied their molecular composition. Starting from the pool of enriched transcripts, we calculated the intersection between the RNAs showing the largest and smallest amounts of protein contacts (eCLIP experiments) [34] against the RNAs showing the largest and smallest amounts of RNA contacts (RISE database) [49]. This approach is useful to reveal the existence of a particular subset specialised in binding specific molecular types. The results are shown in **Supplementary Figure 4**, where we report the intersection of the strongest and poorest protein and RNA binders, comparing them with a control. Despite the vast majority of RNAs does not show significant preference for a certain molecular type, we detected an enrichment in the set of RNAs that binds extensively both proteins and RNAs (**Supplementary Table 3**). Thus, these data confirm that the interactivity of the RNAs enriched in SGs and PBs with boths proteins and RNAs is significantly higher than the other transcripts, supporting their importance in sustaining the network of these biological condensates.

### SGs and PBs protein pairs are enriched in structural disorder

We next investigated the properties of proteins accumulating in SGs and PBs to better understand how they contribute to their interaction network. First, we analyzed how structure affects the formation of protein pairs in these biological condensates in comparison with the rest of the proteins.

We retrieved from BioGRID [50] all binary protein-protein interactions (PPIs) involving proteins located in SGs and PBs (**Materials and Methods** and **Supplementary Table 2)** and, as a control, an equal amount of PPI with interactors that were not found therein (extracted multiple times, **Supplementary Figure 5**). In this analysis, we measured the amount of disorder available using MobiDB (mean disHL disorder score for each pair) [51] (**Figure 3A** and **Supplementary Figure 5** and **6**). In addition, we also measured the amount of disorder of single condensates and non-condensates proteins (**Supplementary Figure 7)**. Both analyses indicated that proteins from PBs and SGs are more disordered than the rest of the proteome.

**Figure 3.**
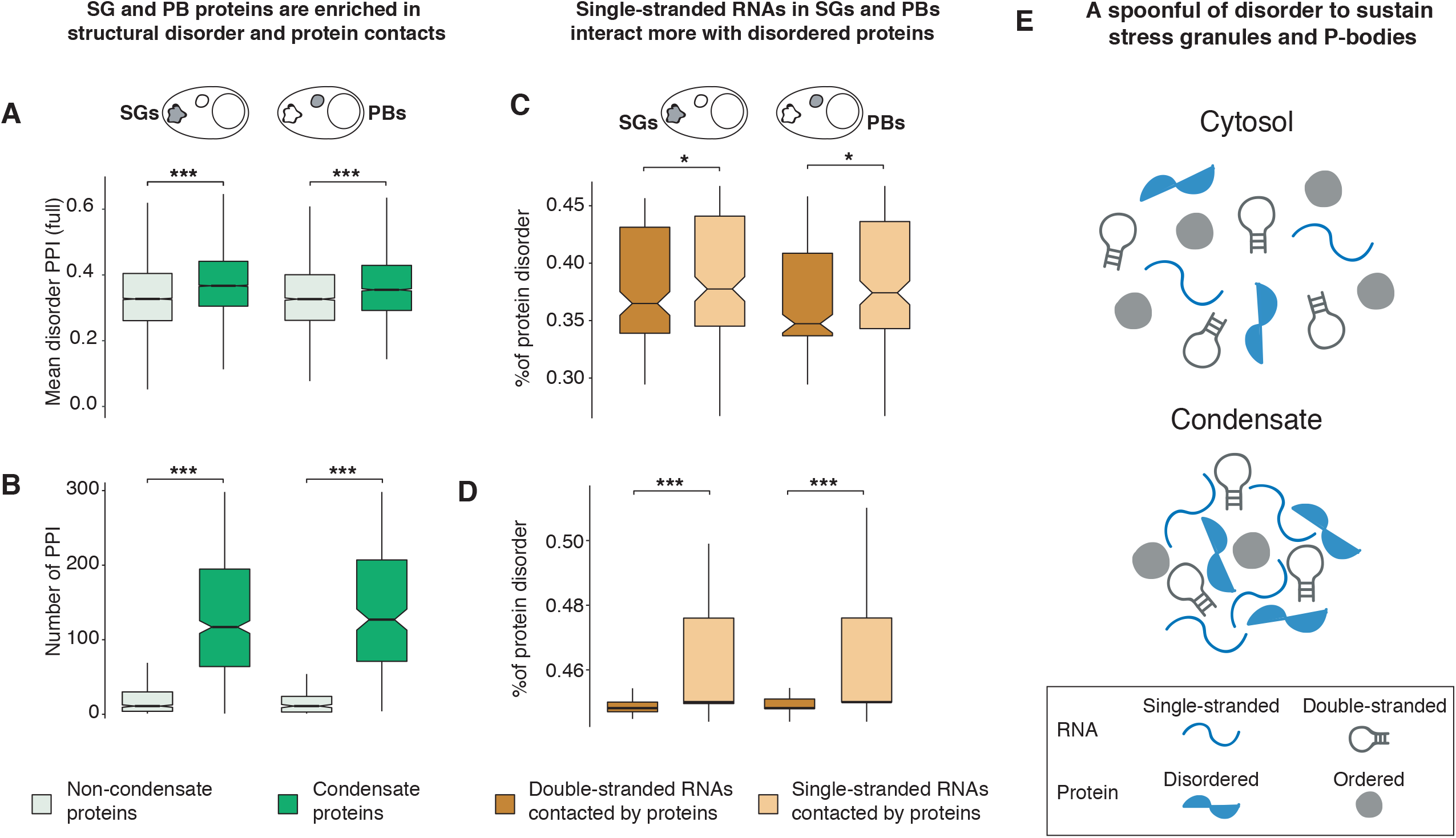
Protein interaction in SGs and PBs is lead by disorder. **A**. Disorder content of protein-protein interactions associated with SGs and PBs (BioGRID database). For each organelle (SG and PB), an equal number of protein pairs (9336 for SG, 3920 for PB) with the non-condensate control is used. The mean disorder content of each pair was retrieved from the MobiDB database (disHL score). Significant differentiation is found (SG p-value <1.05e-157, PB p-value < 2.21e-33, Wilcoxon test). **B**. Number of protein-protein interactions associated with SGs and PBs proteins (BioGRID database). For each organelle (SG and PB), an equal number of proteins (586 for SG, 231 for PB) with the non-condensate control is used. Significant differentiation is found (SG p-value < 3.36e-126, PB p-value < 1.51e-58, Wilcoxon test). C. *cat*RAPID predictions of protein interactions with RNAs most single stranded and double stranded in SGs and PBs and calculation of mean proteins disorder content. An equal amount of 200 transcripts is used in each category (SGs and PBs, most single-stranded and double-stranded RNAs). The mean disorder content of the interacting proteins for each RNA is retrieved from MobiDB (disHL score). Significant differentiation is found (SG p-value < 0.056, PB p-value < 0.067, Wilcoxon test). **D**. Disorder content of eCLIP proteins interacting with SG and PB most single stranded and double stranded RNAs. An equal amount of 200 transcripts is used in each category (SGs and PBs, most single-stranded and double-stranded RNAs). The mean disorder content of the interacting proteins for each RNA is retrieved from MobiDB (disHL score). (SG p-value < 2.89e-17, PB p-value < 4.97e-19, Wilcoxon test). **E**. Graphical representation of interaction patterns found in our analysis. Condensate enriched RNAs and proteins are responsible for the creation of a contact network that involves both other RNAs and proteins. This leads to the hypothesis that granule proteins and enriched RNAs are crossed “hubs” recruiting and sustaining the different components of SGs and PBs. Significance indicated in the plots: ^*^ p-value < 0.1, ^**^ p-value < 0.01 and ^***^ p-value < 0.001.

Thus, in addition to RNAs with increased content of single-stranded regions, our results indicate that SGs and PBs contact networks are enriched in proteins with a lower amount of structure. Since it is known that structural disorder promotes allosteric interactions and favours binding with many protein partners [52], we speculated that SG and PB proteins could have a large number of contacts.

To this aim, we took proteins from SGs and PBs and as a control an equal number of proteins that were not found therein (extracted multiple times, **Supplementary Figure 8**). We then counted how many interactions were reported in BioGRID (**Figure 3B** and **Supplementary Figure 8**) [50]. Our results indicate that SG and PB proteins have a significantly larger number of partners, suggesting that they have a denser contact network than the rest of the proteome. Therefore, we found an equivalence between RNAs and proteins enriched in SGs and PBs, in which both are characterized by a larger number of contacts.

### Disorder proteins in SGs and PBs interact more with linear RNA

Since RNA binding proteins (RBPs) contain disordered regions [53], and SG and PB contact networks are enriched in disordered proteins, we investigated which type of structural properties regulate the interactions between RNAs and proteins. Based on the increased amount of single-stranded regions in enriched RNAs and their capacity to form a larger number of interactions with proteins, we expected an increased amount of disorder in RBPs. To test this hypothesis, we analyzed the least and most structured RNAs (data from DMS measured *in vivo*) [38] in SGs and PBs and measured the disorder content (disHL score from MobiDB) of the interacting proteins [51] (**Materials and Methods**). In this analysis we focused on proteins that bind to RNA as predicted by *cat*RAPID and for which the eCLIP interactome is available [34]. The analysis shows that single-stranded RNAs in SGs and PBs are preferentially contacted by disorder proteins (**Figure 3C)**. The same result was obtained considering interactions from eCLIP experiments, which confirms the validity of our predictions (**Figure 3D**). In the same way, this analysis, carried out only on the RNAs enriched in SGs and PBs, also reproduces this trend (**Supplementary Figure 9)**.

From two independent points of view we arrived at the same conclusion about the organization of molecules contained in SGs and PBs (**Figure 3E**). Enriched RNAs, which are more single-stranded, form a larger number of interactions with RNAs but also have a strong potential to interact with proteins. Disordered proteins are enriched in SGs and PBs, have a larger number of PPIs, but also can form more contacts with single-stranded RNAs. So, the two molecular sets that we detected as the most interacting, RNAs and proteins, are both depleted in structure, and form strong interactions between them. This finding indicates that proteins and RNAs in SGs and PBs act together as “hubs” that recruit and sustain the different components of the assemblies (**Figure 3E**).

## DISCUSSION

We previously observed that RNAs enriched in double-stranded regions attract a large number of proteins [19,20]. The origin of this trend, also identified in SG and PB analyses, is that double-stranded regions favor stable interactions with proteins by reducing the intrinsic flexibility of polynucleotide chains [19,20]. While for each amino acid residue there are two torsional degrees of freedom, RNA conformational space is greater -for each nucleotide residue there are seven independent torsion angles.

Here, we report the novel result that RNAs enriched in SGs and PBs contain single-stranded regions that contact specific elements of their binding partners. Since recent reports indicate that single-stranded RNAs have strong ability to act as scaffolds of SGs and PBs [13,30], we primarily focused our analyses on their interactions with proteins and RNAs.

We first found that RNAs enriched in single-stranded regions are prone to engage in RNA-RNA contacts. This result is not unexpected since single-stranded transcripts are able to base-pair [48,54] and, by doing so, establish a network of stable interactions. We note that the analysis of RNA-RNA interactions does not take into account the cellular context in which SGs and PBs are formed, thus our results are compatible with a scenario in which specific transcripts are highly prone to interact to quickly promote molecular condensation [13].

In parallel, the analysis of the SGs and PBs protein interaction networks revealed that proteins enriched in disordered elements form a larger number of contacts with other proteins. This result is very well in line with recent reports indicating that unstructured regions modulate the formation of phase separated assemblies [31]. Indeed, phase separation is a widespread phenomenon in the cell

[55] and disordered interactions greatly contribute to the assembly formation [56]. By reporting that single-stranded RNAs preferably contact disorder proteins, we extended the concept of “fuzziness” to RNA molecules. Our work leads to the intriguing result that the two molecular sets identified as the most interacting in the proteome and transcriptome are both depleted in structure and bind one to the other. Thus, specific elements in proteins and RNAs have the ability to recruit and sustain all the components of SGs and PBs (**Figure 3E**).

In conclusion, our work suggests that there is not only great diversity in the interaction partners (RNA-RNA, protein-protein, and RNA-protein) but also in their binding modes [57,58]. In this complex scheme, RNA ability to induce phase separation can have an impact on both ordered and disordered proteins: while structural elements can irreversibly sequester globular proteins [12], disordered regions dynamically favor phase separation [59]. Our study shows that the inner architectures of SGs and PBs are intrinsically governed by RNAs and proteins with an increased amount of structurally disordered domains. Thanks to the dynamicity of these regions, protein-RNA complexes are able to assemble and disassemble without the need of strong efforts by the cell. RNA-RNA interactions, at present poorly investigated, are expected to greatly contribute to establishing molecular associations within SGs and PBs [13]. Indeed, RNA molecules are versatile platforms [1,40] capable of interacting with all other molecules [19], thus promoting the efficient coordination of transcriptional and post-transcriptional layers of regulation [20].

## MATERIALS AND METHODS

### SG and PB transcriptomes

SG transcriptome was collected from Khong. *et al*. [27]. The data was generated through RNA-sequencing (RNA-seq) analysis of purified SG cores and single-molecule fluorescence *in situ* hybridization (smFISH) validation. The PB transcriptome was retrieved from Hubstenberger. *et al*. [30], in which a fluorescence-activated particle sorting (FAPS) method was used to purify cytosolic PBs from human epithelial cells. In our statistical analysis, we applied filtering and retained only RNAs with an experimental p-value < 0.01. Within the transcriptome, we distinguished two subsets of transcripts depending on their abundance with respect to the cell transcriptome: enriched (fold-change >=2) and depleted (fold-change<=0.5).

### SG and PB proteomes

SG proteome data was retrieved from experiments in various stress conditions and different cell types [7,9,60] for a total of 632 proteins. The first dataset was obtained purifying SG cores from Sodium Arsenite (NaAsO_2_) stressed U-2 OS cells using a series of differential centrifugations and then affinity purification of GFP-G3BP. The second dataset was obtained using a combination of ascorbate peroxidase (APEX)-mediated *in vivo* proximity labeling with quantitative mass spectrometry (MS) and an RBP-focused immunofluorescence (IF) to identify SG proteins in neuronal and non-neuronal cells and under different types of stress conditions (heat shock, ER stress and oxidative stress). The third dataset employs systematic in vivo proximity-dependent biotinylation (BioID) analysis to identify core components of SGs and PBs. PB proteome data was retrieved combining two studies [30,53] for a total of 259 proteins. In the first study, a fluorescence-activated particle sorting (FAPS) method was developed to purify cytosolic PBs from human epithelial cells, while the second dataset is the one mentioned before, which identified core proteins for both PB and SG using BioID analysis.

### RNA secondary structure prediction

We predicted the secondary structure of transcripts using *CROSS* (Computational Recognition of Secondary Structure) [35]. The algorithm predicts the structural profile (single- and double-stranded state) at single-nucleotide resolution using sequence information only and without sequence length restrictions (scores > 0 indicate double stranded regions). The obtained scores are then averaged to obtain a secondary structure propensity score for each transcript.

### RNA secondary structure measured by DMS

Data on RNA structural content measured by dimethyl sulfate modification (DMS) *in vitro* and *in vivo* conditions were retrieved from Rouskin *et al*. [38]. The number of reads of each transcript was normalized to the highest value (as in the original publication) and averaged.

### RNA secondary structure measured by PARS

To profile the secondary structure of human transcripts, we used Parallel Analysis of RNA Structure (PARS) data [36]. To measure PARS structural content for each transcript, we computed the fraction of double-stranded regions over the entire sequence. Given the stepwise function □ (*x*)□=□1 for *x*□>□0 and LJ(*x*)□=□0 otherwise, we computed the fraction of structured domains as:

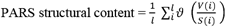

where *V*(*i*) and *S*(*i*) are the number of double- and single-stranded reads.

To measure the secondary structure content of the human transcripts 5′- and 3′-UTR and CDS, we retrieved the corresponding locations of the 5′- and 3′-UTR from Ensembl database and repeated the same procedure described above simply considering only the corresponding part of the sequence.

### Protein-RNA interaction prediction

Predicted interactions with human proteins were retrieved from RNAct [47], a database of protein-RNA interactions calculated using *cat*RAPID *omics* [61], an algorithm to estimate the binding propensity of protein-RNA pairs by combining secondary structure, hydrogen bonding and van der Waals contributions [44]. As reported in the analysis of about half a million of experimentally validated human interactions [47], the algorithm is able to separate interacting vs non-interacting pairs with an area under the ROC curve of 0.78 [62]. The output is filtered according to the Z-score column, which is the interaction propensity normalised by the mean and standard deviation calculated over the reference RBP set. For our analysis, we considered only predicted interactions with a Z-score > 1.

### Experimental data on Protein-RNA interactions

RNA interactions for 151 RBPs were retrieved from eCLIP experiments [63] performed in K562 and HepG2 cell lines. In order to measure the fraction of protein binders for each transcript, we applied stringent cut-offs [-log_10_(*p*-value) >5 and -log_2_(fold_enrichment) >3] as in previous work [63]. Furthermore, in case of interactions established in one cell line, only interactions seen in 2 replicates were retained, while in case of two cell lines, only interactions seen in at least 3 out of 4 replicates were retained.

### RNA-RNA interactions predictions

RNA-RNA interactions were predicted using the stand-alone IntaRNA software [48], a program for the fast and accurate prediction of interactions between two RNA molecules. It has been designed to predict mRNA target sites for given non-coding RNAs, such as eukaryotic microRNAs (miRNAs) or bacterial small RNAs (sRNAs), but it can be used to predict other types of RNA-RNA interactions. For each predicted RNA-RNA interaction we retrieved the most optimal one and considered the associated interaction energy.

### Experimental data on RNA-RNA interactions

Information about human RNA-RNA interactions were retrieved from RNA Interactome from Sequencing Experiments (RISE) database [49]. RISE is a comprehensive repository of RNA-RNA interactions that mainly come from transcriptome-wide sequencing-based experiments such as PARIS, SPLASH, LIGRseq, and MARIO, and targeted studies like RIAseq, RAP-RNA, and CLASH. Currently it hosts 328,811 RNA-RNA interactions mainly coming from three species (human, mouse, yeast). Human RNA-RNA interactions were filtered, and we retrieved only those in which both partners had an available Ensembl ID.

### Experimental data on protein-protein interactions

We used BioGRID (version 4.2.193) for experimental data on protein-protein interactions data [50]. BioGRID is a biomedical interaction repository with data compiled through comprehensive curation efforts, and it contains protein and genetic interactions, chemical interactions and post translational modifications from major model organism species. We used BioGRID to retrieve protein-protein interactions involving condensates proteins against a control. To further strengthen our results, our analyses were done considering both the entire available human BioGRID interactome and physical interactions.

### Protein disorder information

Information about human protein disorder predictions were retrieved from MobiDB database (version 4.0) [51], that contains several data resources and features for protein disorder. Structural and functional properties of disordered regions are based on third party databases and a set of prediction methods, which are assembled to provide a comprehensive view of properties of disordered regions at the residue level. From the whole set of predictive methods, we selected scores obtained with DisEMBL tool with hot loops threshold (DisHL), developed for the prediction of loops with a high degree of mobility, considered important for the definition of protein disorder [64].

### Statistical analysis

To assess the significance of the different trends throughout the analysis, we used the Wilcoxon rank sum test (two-sided). It is a non-parametric test that can be used to compare two independent groups of samples. In order to have analysis with balanced groups, for each comparison performed in our study we used the same number of RNAs/proteins for each category, except when stated otherwise.

## ACKNOWLEDGMENTS

The authors would like to thank all the members of Tartaglia’s lab at the CRG and IIT and especially Alexandros Armaos.

