## Supplementary Materials for "Interplay between disordered regions in RNAs and proteins modulates interactions within stress granules and processing bodies"

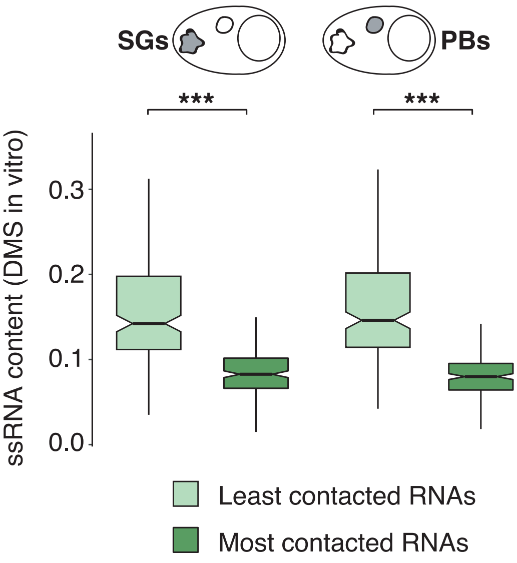


**Supp. Figure 1.** Single stranded content (dimethyl sulfate modification, DMS, measured *in vitro*) for RNAs most and least contacted by proteins in SGs and PBs [1]. The RNAs are categorized in least or most contacted depending on the amount of protein interactions detected by eCLIP [2]. An equal amount of 200 transcripts is used in each category (SGs and PBs, least and most contacted RNAs). Significant differentiation is found (SG p-value < 4.34e-38, PB p-value < 1.01e-42, Wilcoxon test)

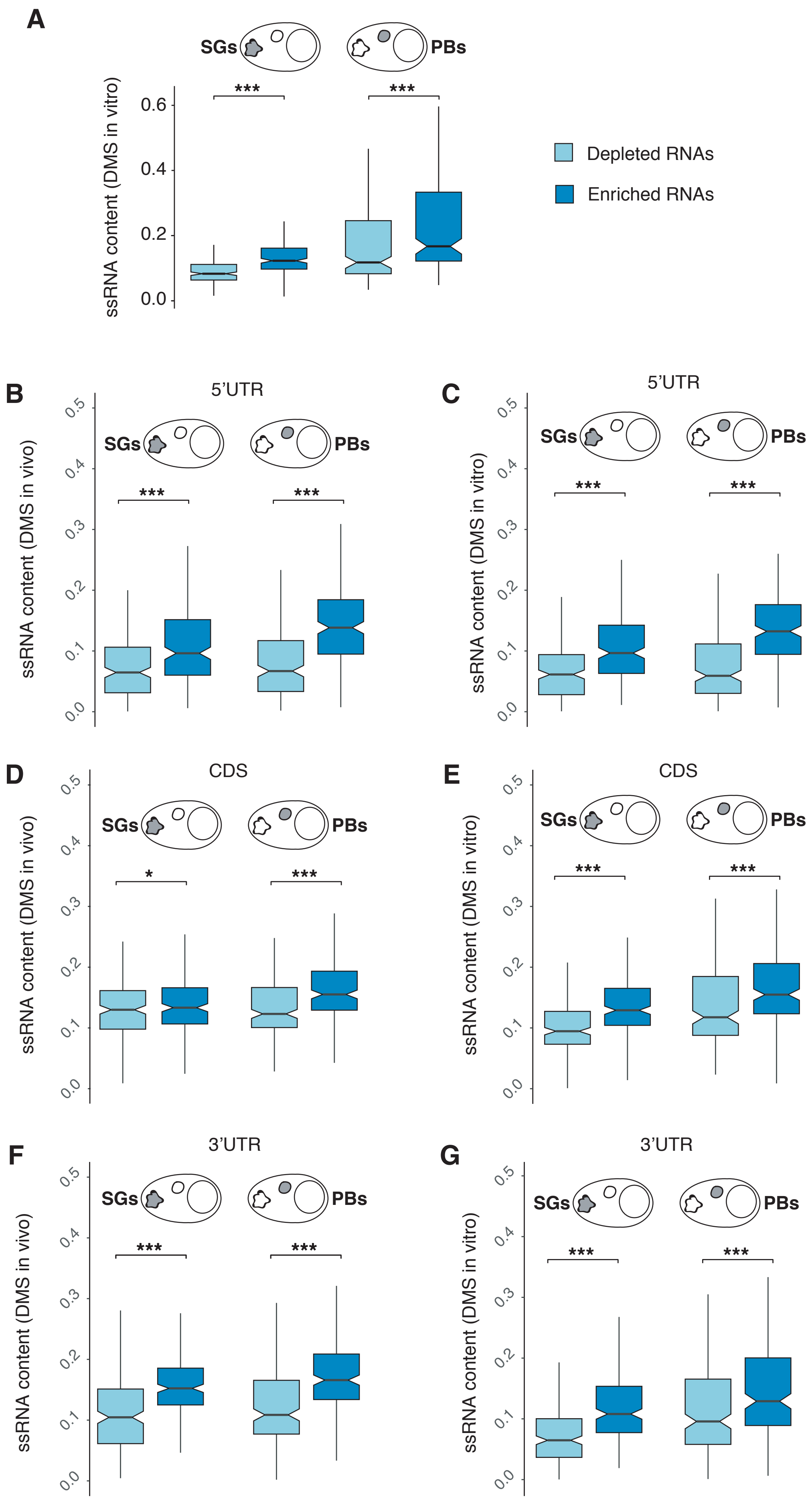
A
SGs
PBs
***
***
ssRNA content (DMS in vitro)
0.0
0.2
0.4
0.6
Depleted RNAs
Enriched RNAs
B
5’UTR
SGs
PBs
***
***
ssRNA content (DMS in vivo)
0.0
0.1
0.2
0.3
0.4
0.5
C
5’UTR
SGs
PBs
***
***
ssRNA content (DMS in vitro)
D
CDS
SGs
PBs
*
***
ssRNA content (DMS in vivo)
E
CDS
SGs
PBs
***
***
ssRNA content (DMS in vitro)
F
3’UTR
SGs
PBs
***
***
ssRNA content (DMS in vivo)
G
3’UTR
SGs
PBs
***
***
ssRNA content (DMS in vitro)

**Supp. Figure 2.** Single stranded content of RNA regions (ssRNA). An equal amount of 200 transcripts is used for each category (SGs and PBs, depleted and enriched RNAs). **A.** Single stranded content (DMS, measured *in vitro*) for RNAs enriched or depleted in SGs and PBs [1]. Significant differentiation is found (SG p-value < 7.17e-72, PB p-value < 2.36e-76, Wilcoxon test). **B.** Single stranded content (DMS, measured *in vivo*) for RNAs enriched or depleted in SGs and PBs, considering the 5'UTR part of the sequence [1]. Significant differentiation is found (SG p-value < 8.65e−09, PB p-value < 1.16e−21, Wilcoxon test). **C.** Single stranded content (DMS, measured *in vitro*) for RNAs enriched or depleted in SGs and PBs, considering the 5'UTR part of the sequence [1]. Significant differentiation is found (SG p-value < 6.93e−12, PB p-value < 2.64e−15, Wilcoxon test). **D.** Single stranded content (DMS, measured *in vivo*) for RNAs enriched or depleted in SGs and PBs, considering the CDS part of the sequence [1]. Significant differentiation is found (SG p-value < 0.059, PB p-value < 2.42e−08, Wilcoxon test). **E.** Single stranded content (DMS, measured *in vitro*) for RNAs enriched or depleted in SGs and PBs, considering the CDS part of the sequence [1]. Significant differentiation is found (SG p-value < 5.56e−14, PB p-value < 5.64e−08, Wilcoxon test). **F.** Single stranded content (DMS, measured *in vivo*) for RNAs enriched or depleted in SGs and PBs, considering the 3'UTR part of the sequence [1]. Significant differentiation is found (SG p-value < 8.80e−17, PB p-value < 3.53e−15, Wilcoxon test). **G.** Single stranded content (DMS, measured *in vitro*) for RNAs enriched or depleted in SGs and PBs, considering the 3'UTR part of the sequence [1]. Significant differentiation is found (SG p-value < 3.80e−15; PB p-value < 1.07e−06, Wilcoxon test).

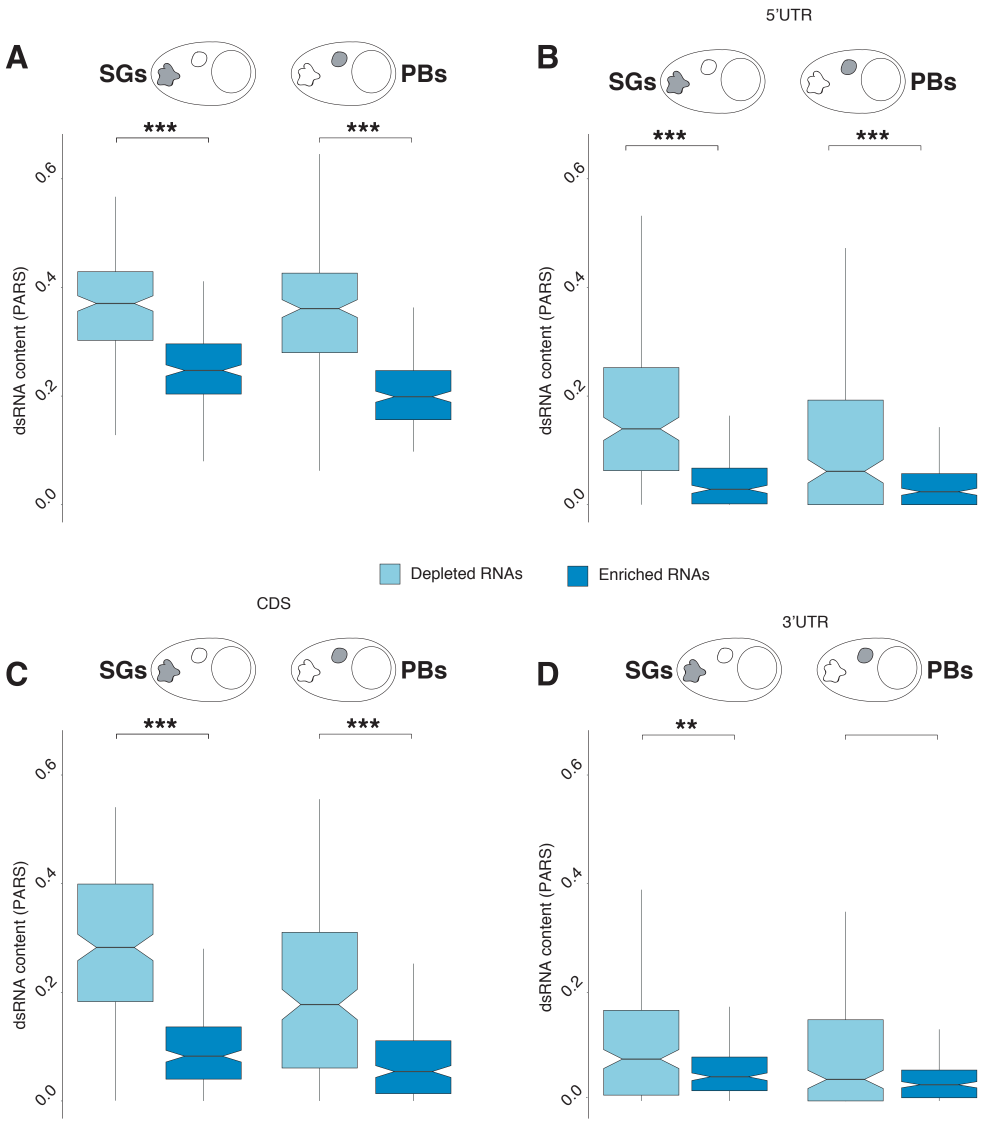


**Supp. Figure 3.** Double stranded content of RNA regions (dsRNA). An equal amount of 200 transcripts is used for each category (SGs and PBs, depleted and enriched RNAs). **A.** Double stranded content (PARS technique) for RNAs enriched or depleted in SGs and PBs [3]. Significant differentiation is found (SG p-value < 7.92e−27, PB p-value < 8.34e-31, Wilcoxon test). **B.** Double stranded content (PARS technique) for RNAs enriched or depleted in SGs and PBs, considering the 5'UTR part of the sequence [3]. Significant differentiation is found. (SG p-value < 4.22e-26, PB p-value < 2.08e-06, Wilcoxon test). **C.** Double stranded content (PARS technique) for RNAs enriched or depleted in SGs and PBs, considering the CDS part of the sequence [3]. Significant differentiation is found (SG p-value < 2.02e-35, PB p-value < 8.16e-15, Wilcoxon test). **D.** Double stranded content (PARS technique) for RNAs enriched or depleted in SGs and PBs, considering the 3'UTR part of the sequence [3]. Significant differentiation is found. (SG p-value < 0.002; PB p-value < 0.144, Wilcoxon test).

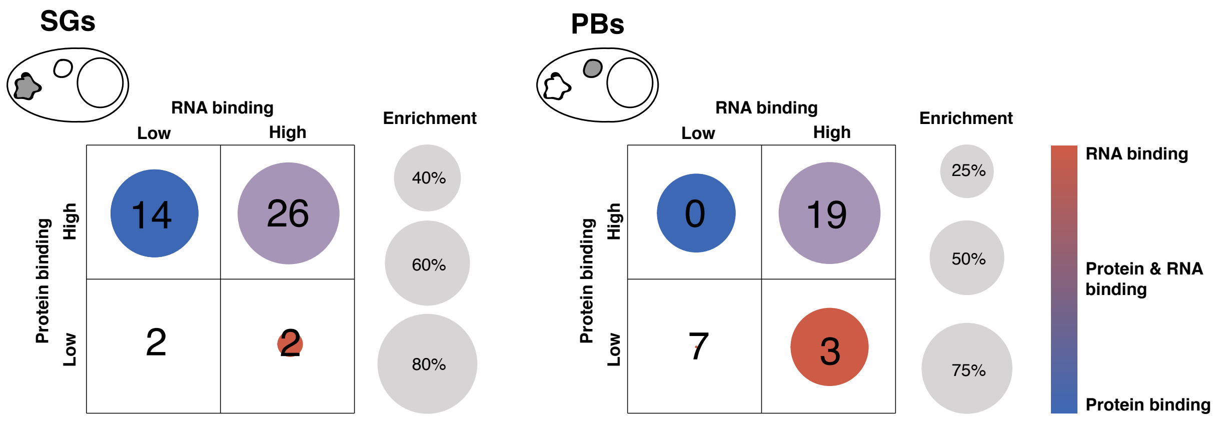


**Supp. Figure 4.** Composition of enriched RNAs in SGs and PBs. The numbers in the table show the intersection between the RNAs most and least contacted by proteins (eCLIP experiments) [2] against the RNAs most and least contacted by RNAs (RISE datasets) [4] of RNAs enriched in SGs and PBs. An equal amount of 100 transcripts is used in each category (protein-RNA and RNA-RNA interactions). 1000 random extractions of non-enriched human RNAs sets were used as control (same size as the target). The circles size shows the probability of the target to be larger than the control.

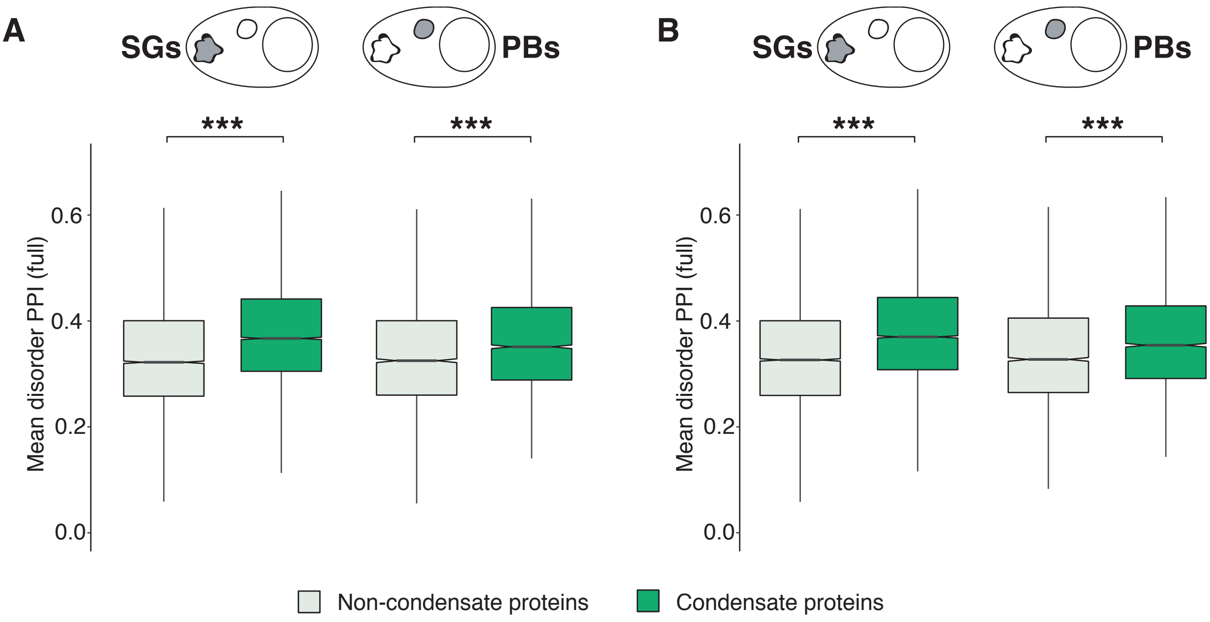


**Supp. Figure 5. A.** Disorder content of protein-protein interactions associated with SGs and PBs (BioGRID database) [5] calculated on the second random extraction for the control. For each organelle (SG and PB), an equal number of protein pairs (9336 for SG, 3920 for PB) with the non-condensate control is used. The mean disorder content of each pair was retrieved from MobiDB database (disHL score) [6]. Significant differentiation is found (SG p-value < 3.61e-191, PB p-value < 9.73e-30, Wilcoxon test). **B.** Disorder content [6] of protein-protein interactions associated with SGs and PBs (BioGRID database) [5] calculated on the third random extraction for the control. The number of protein pairs follow the definition given in panel **A.** Significant differentiation is found (SG p-value < 1.95e-190, PB p-value < 1.57e-26, Wilcoxon test).

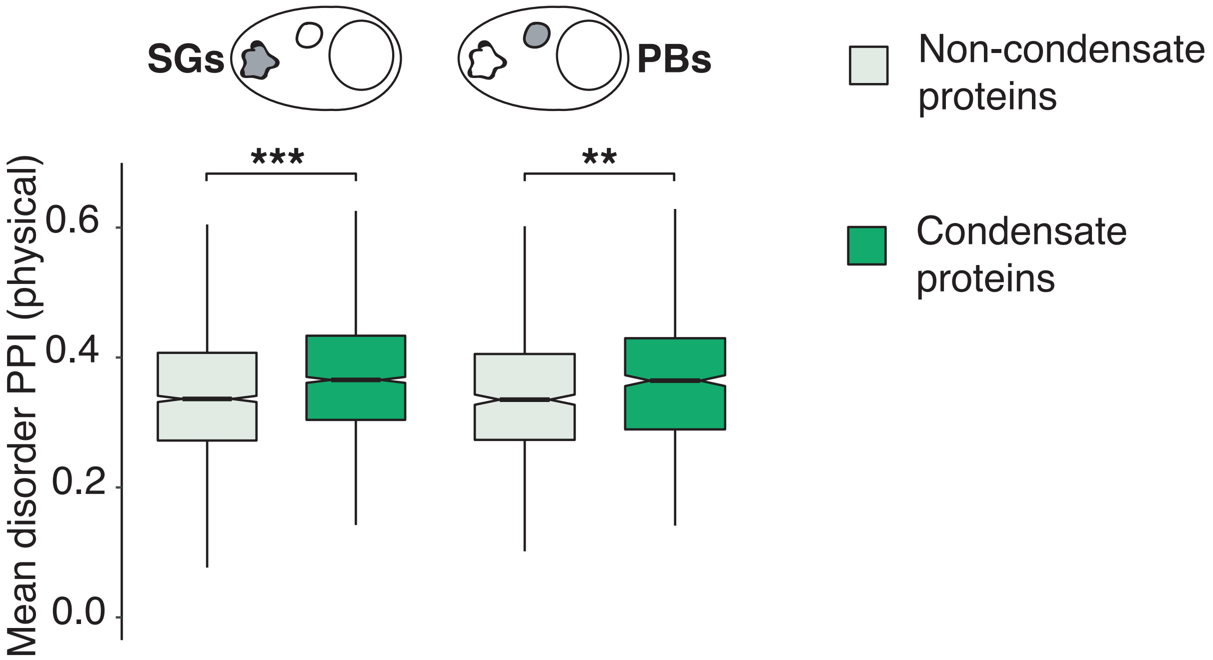


**Supp. Figure 6.** Disorder content [6] of protein-protein interactions associated with SGs and PBs (BioGRID database, physical interactions only) [5]. For each organelle (SG and PB), an equal number of protein pairs (1997 for SG, 704 for PB) with the non-condensate control is used. The mean disorder content of each pair was retrieved from MobiDB database (disHL score) [6]. Significant differentiation is found (SG p-value < 2.23e-19, PB p-value < 0.002, Wilcoxon test).

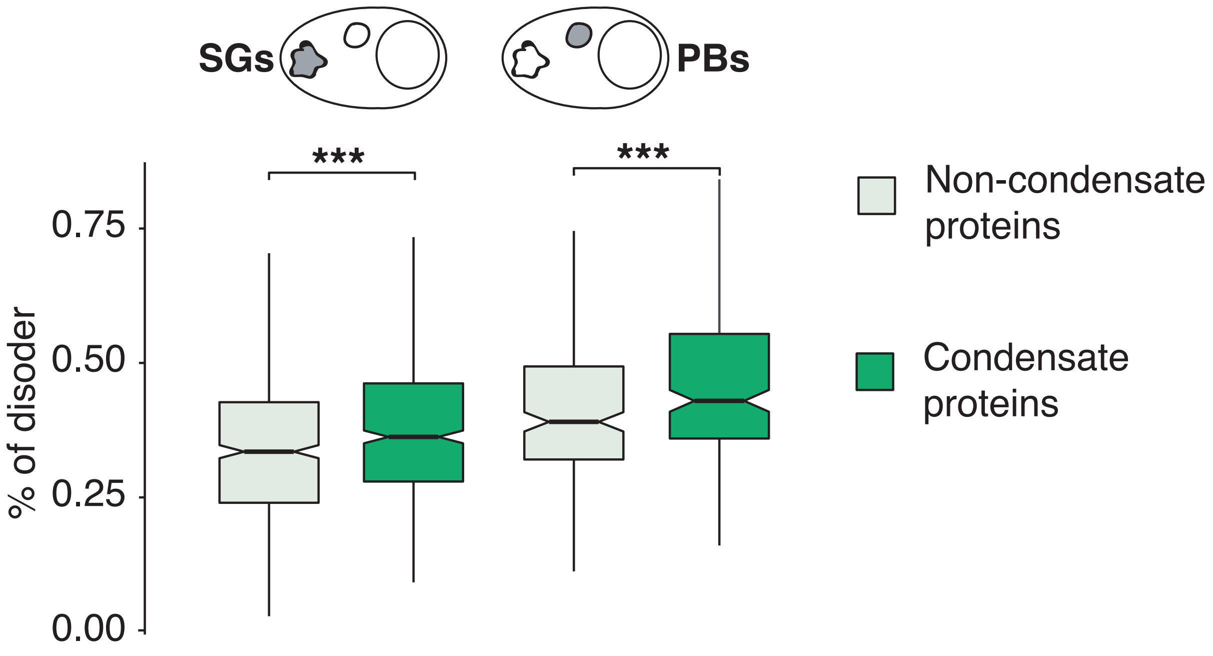


**Supp. Figure 7.** Disorder content [6] of condensate and non-condensate proteins. For each organelle (SG and PB), an equal number of proteins (586 for SG, 231 for PB) with the non-condensate control is used. The disorder content is retrieved from the MobiDB database (disHL score) [6]. Significant differentiation is found (SG p-value < 5.49e-05, PB p-value < 2e-04, Wilcoxon test).

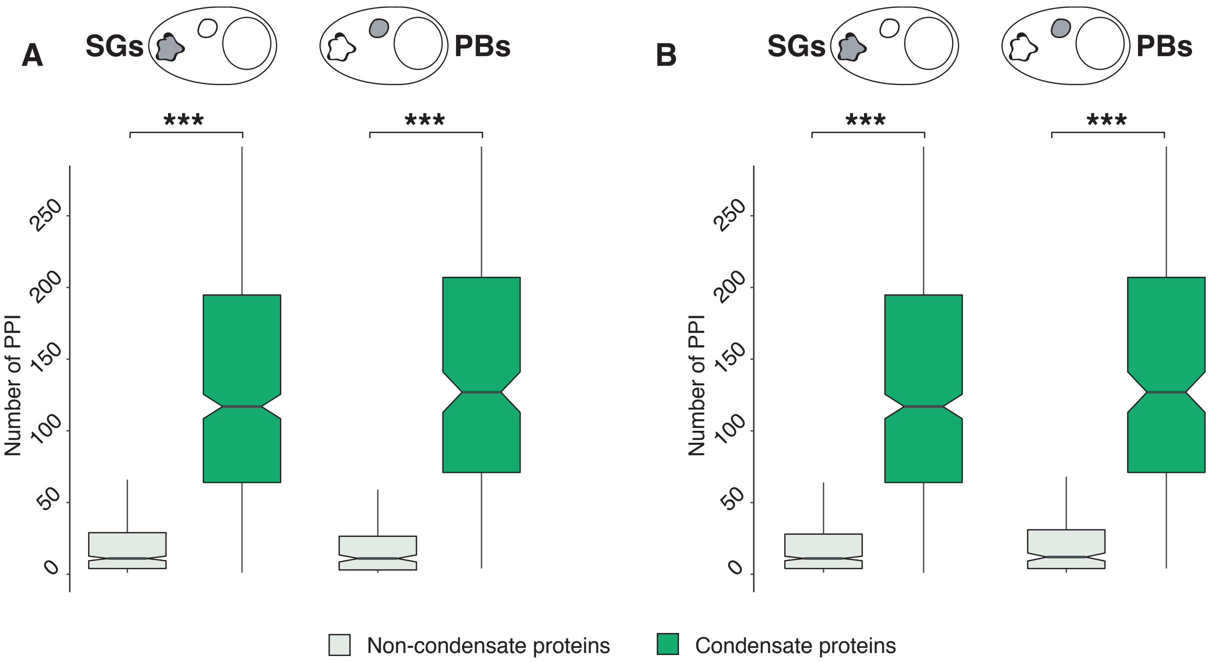


**Supp. Figure 8. A.** Number of protein-protein interactions associated with SGs and PBs proteins (BioGRID database) [5] with the second random extraction of the control. For each organelle (SG and PB), an equal number of proteins (586 for SG, 231 for PB) with the non-condensate control is used. Significant differentiation is found (SG p-value < 1.11e-128, PB p-value < 5.73e-59, Wilcoxon test). **B.** Number of protein-protein interactions associated with SGs and PBs proteins (BioGRID database) [5] with the third random extraction of the control. Proteins classes follow the definition given in panel **A.** Significant differentiation is found (SG p-value < 7.21e-129, PB p-value < 1.10e-53, Wilcoxon test).

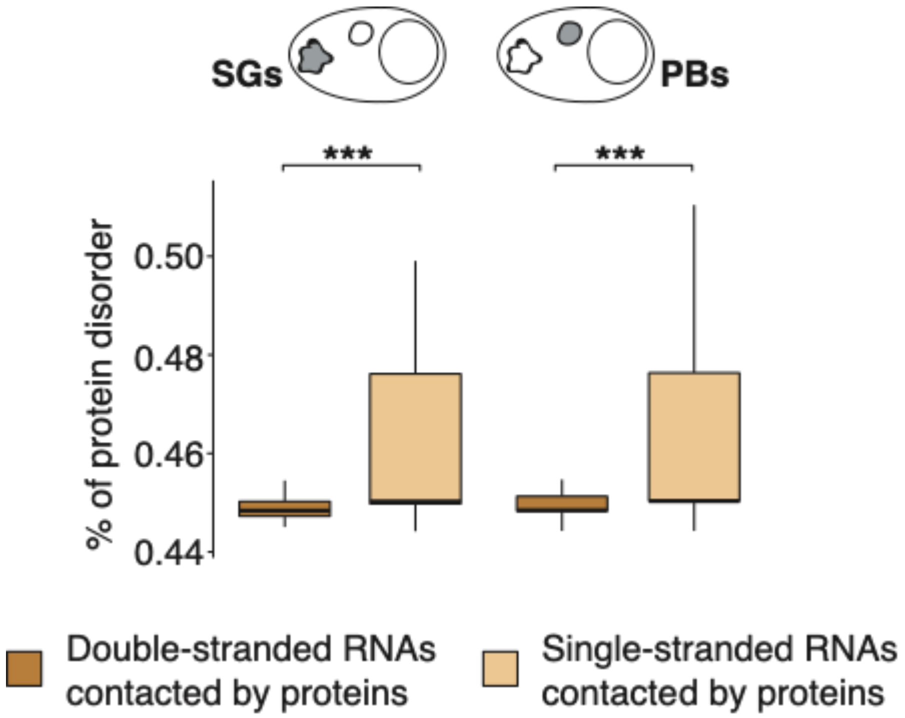


**Supp. Figure 9.** Disorder content [6] of eCLIP proteins [2] interacting most single stranded and double stranded RNAs that are enriched in SGs and PBs [1]. An equal amount of 200 transcripts is used in each category (SGs and PBs, enriched RNAs most single-stranded and double-stranded RNAs). The mean disorder content of the interacting proteins for each RNA is retrieved from MobiDB (disHL score) [6]. Significant differentiation is found (SG p-value < 1.02e-27, PB p-value < 2.09e-21, Wilcoxon test).

**Supplementary Table 1.** SGs and PBs transcriptomes used in the analysis. All the RNAs listed in the table are coding transcripts with an associated P-value from the original experiment < 0.01.

**Supplementary Table 2.** SG and PB proteomes, retrieved combining different experimental datasets obtained under various stress conditions and in different cell types and collected with diverse purification techniques, for a total of 632 proteins in SG and 259 in PB.

**Supplementary Table 3.** SG and PB RNAs most contacted by both RNAs (RISE) [4] and proteins (eCLIP) [2].
